# RAPTORGraph: graph-based pathway modeling for causal discovery in single-cell perturbations

**DOI:** 10.64898/2026.11.03.763980

**Authors:** Yeremia Gunawan Adhisantoso, Stephanie Kristin Schröder, Maximilian Greß, Mikel Hernaez, Jan Voges

**Author notes:** Corresponding authors: Mikel Hernaez and Jan Voges.

## Abstract

Experiments involving the perturbation of individual cells are central to understanding cellular mechanisms and can accelerate drug discovery. Causal representation learning allows us to uncover the latent factors that regulate biological systems and predict the impact of novel perturbations. Unfortunately, existing methods fail to address intervention spillover in a closed-world setting where intervention targets are known *a priori*, such as in Perturb-seq experiments, due to their reliance on dense encoders. Furthermore, incorporating curated biological pathways into the model imposes a confirmatory bias, forcing it to explain the data through preexisting pathways and reducing the set of hypotheses the model can explore, while discarding novel signals that lie outside the annotated pathways. In this work, we introduce RAPTORGraph, a *β*-variational autoencoder with a GraphPathway encoder that explicitly models complex gene-to-gene interactions within learned pathways. Moreover, our model’s preconditioning isolates the influence of perturbed genes, yielding clean, single-node latent interventions required for identifiable causal discovery and eliminating spillover. Finally, we train the model on data preprocessed with optimal-transport alignment, which guarantees a well-defined mapping between control and perturbed samples and further stabilizes the learned latent representations. We demonstrate that RAPTORGraph improves state-of-the-art performance on downstream analyses of unseen perturbations, such as non-additive interactions, while performing competitively on objective metrics, such as mean squared error and maximum mean discrepancy. The source code is available at https://github.com/sXperfect/RAPTORGraph.

**Author summary:** Understanding how genes work together can help explain changes in cell behavior and guide the search for treatments. Yet testing every possible combination of genetic changes is impractical. We developed a computer model to predict how individual cells respond to changes that have not been tested experimentally. We address two problems: existing models can obscure how a change leads to a predicted outcome and overlook differences between individual cells. In our model, each genetic change enters through a single starting point, and its effects can spread through a network learned from the data. This allows us to examine how a predicted response develops. We also match cells from separate groups with and without genetic changes, helping our model retain the diversity of cellular responses. In tests on two datasets, we improved predictions of responses to genetic perturbations. On the Norman et al. dataset, we also improved predictions of responses to combined genetic changes whose effects could not be explained by adding the effects of the individual changes. We also used the model to suggest genetic changes that could explain an observed cellular response. Our approach could help researchers choose informative experiments and investigate how genes jointly shape cell behavior.

## Introduction

Understanding how cells respond to genetic perturbations, such as those induced by CRISPR screens [1], is fundamental to advancing our understanding of basic biology and therapeutic discovery [2–4]. However, the combinatorial explosion across *∼*20,000 protein-coding genes makes exhaustive perturbation experiments infeasible [4], motivating computational models that predict cellular responses to unseen perturbations or combinations thereof.

Existing approaches to predict the transcriptional outcome following a perturbation face a fundamental challenge: generative quality comes at the cost of causal interpretability. Models such as GEARS [5] and scGPT [6] have demonstrated the ability to predict transcriptional responses to genetic perturbations, including previously unseen perturbations, although their predictive performance varies across datasets and evaluation metrics; however, they act as “black boxes” providing little mechanistic insight into how perturbations affect cellular systems. This lack of interpretability prevents researchers from understanding why certain combinations of perturbations produce synergistic effects, failing to provide testable biological hypotheses.

As such, computational biology demands models that provide mechanistic insights into cellular function [7], making causal representation learning (CRL) [8] a promising approach for mapping genes to biological processes. In particular, recent CRL approaches such as DiscrepancyVAE (dVAE) [9] or SENA [10] aim at recovering the underlying causal generative factors and their interactions, enabling counterfactual reasoning. However, these CRL approaches suffer from fundamental architectural limitations that prevent them from fulfilling their theoretical causal identifiability guarantees: the use of a dense interventional encoder inherently means that every gene, including those intervened upon, can influence all latent variables.

We term this phenomenon *intervention spillover*, as it renders the task of identifying a unique latent target for an intervention ill-posed. Consequently, any mechanism designed to apply the intervention is forced to converge to a suboptimal solution, because it cannot perform the clean, single-node manipulation required for valid causal inference [11, 12].

Further, the destructive nature of single-cell RNA sequencing (scRNA-seq) makes it impossible to observe the same cell before and after a genetic intervention, complicating the prediction of cell-specific responses. Existing methods address this issue by randomly pairing control and perturbed cells. Since the model must minimize error across all possible arbitrary pairings, it is incentivized to learn the population-mean response. We refer to this phenomenon as *mean collapse*, as this regression to the mean erodes cellular diversity.

We present **R**esponse **A**nalysis of **P**erturbed **T**ranscript**O**mes using Inte**R**pretable **Graph** (RAPTORGraph), a novel framework that addresses causal identifiability challenges in perturbation prediction through structured encoder design. Our first key innovation is the GraphPathway layer, a novel architecture that uses preconditioning to enforce sparse mappings from genes to learned latent factors. This design enables the model to perform the clean, single-node manipulations required for valid causal inference, directly maintaining interpretability. This innovation is complemented by a novel optimal transport (OT) preprocessing step, which mitigates the distinct problem of mean collapse. We benchmark the generative quality of RAPTORGraph on the Norman [4] Perturb-seq dataset and the Replogle [13] CRISPRi dataset. For the Norman data, we further evaluate the accuracy of genetic interaction in perturbation combinations, and the biological interpretability of the inferred latent factors. Specifically, when applied to scRNA-seq data from a K562 cell line, RAPTORGraph uncovered distinct, biologically accurate programs, such as the differentiation-induced cell cycle arrest driven by EGR1 and stress-induced growth arrest mediated by JUN. We demonstrate strong performance alongside interpretability, which addresses key gaps not filled by current approaches.

## Background

### In silico high-throughput screening models

Accurate *in silico* prediction of perturbation effects remains critical to guide experimental perturbation assays, as it would allow us to predict, for instance, how cells respond to environmental stress [14]. However, a key limitation in learning perturbation responses lies in the destructive nature of single-cell transcriptomic assays, which prevents measuring the same cell before and after a perturbation. While paired measurements are infeasible, we do have access to independent single-cell observations from control and perturbed cells.

Recent works have demonstrated the potential of representation learning in leveraging these perturbation data to predict transcriptional outcomes of novel combinations of perturbations. For example, Kamimoto et al. [15] relied on inferring (linear) transcriptional relationships between genes in the form of a gene regulatory network to infer the effect of unseen perturbations. Shortly after, the Compositional Perturbation Autoencoder (CPA) [16] predicted combinations of perturbations by vector arithmetic in the latent space generated by a Variational Autoencoder (VAE). In parallel, GEARS [5] leveraged Graph Neural Networks (GNNs) to incorporate prior knowledge of gene–gene relationships into the model architecture, thereby predicting unseen genetic interventions and their combinations. More recently, CellOT [17] proposed to directly learn optimal transport maps between control and perturbed cell states, thus accounting for heterogeneous subpopulation structures in the data. Additional efforts, such as scGen [18] and scGPT [6], have explored deep generative approaches for this task.

Even though all the outlined approaches tackle the common problem of predicting the transcriptional consequences of perturbations, their non-causal “black box” nature hampers their capability for uncovering the causal mechanisms driving observed transcriptional changes. This poses a significant limitation in, for example, cellular engineering, where understanding the *why* behind perturbation effects is as critical as predicting the *what*. To address this challenge, recently, dVAE [9] learns causal latent factors without an explicit semantic meaning, resulting in causally valid but biologically uninterpretable latent representations. SENA [10], on the other hand, explicitly models (a set of) biological processes as latent causal variables, promising both predictive accuracy and mechanistic interpretability.

### Structural causal models for latent discovery

We model the underlying data-generating process using a structural causal model (SCM) [19]. We assume that observed high-dimensional samples **x** *∈* R^*n*^ (e.g., gene expression) are generated by a set of unobserved, low-dimensional latent causal variables **u** *∈* R^*d*^, where *d ≪ n*. The SCM defines the relationships between these variables via a set of structural assignments:

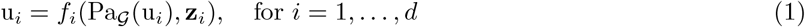

where *f*_*i*_ is the structural function, Pa_*G*_ (u_*i*_) are its direct causal parents in a graph *G*, and **z**_*i*_ is an independent exogenous noise term. The complete set of these equations induces a directed acyclic graph (DAG) *G*, which implies that the joint probability distribution over **u** factorizes as:

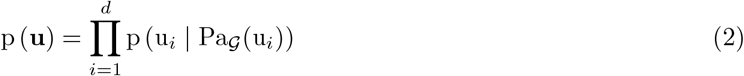

However, these latent causal variables are not directly accessible. Instead, we only observe the high-dimensional data **x**, which are generated via a true, unknown mixing function *g*^*∗*^:

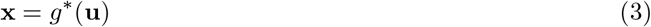

### Identifiability from targeted interventions

The central challenge of CRL is *identifiability* : learning an encoder function *f*_*θ*_ that can uniquely recover the true latents **u** from only observations **x**. However, using observational data alone, this goal is theoretically intractable [19]. In this sense, Khemakhem et al. [20] showed that different parameter configurations of a VAE can generate equivalent marginal distributions p (**x**), leaving the learned latent representations non-unique. In the causal setting, this problem is compounded, as multiple distinct causal structures can generate identical observational distributions. This means that learning interpretable causal mechanisms from observational data alone is ill-posed.

To overcome this fundamental limitation, modern CRL relies on access to interventional data. The key insight is that interventions create unique statistical signatures that break the symmetries present in observational data (Fig. 1). As formally established in recent work [8], achieving identifiability depends on a set of key requirements.

**Fig 1.**
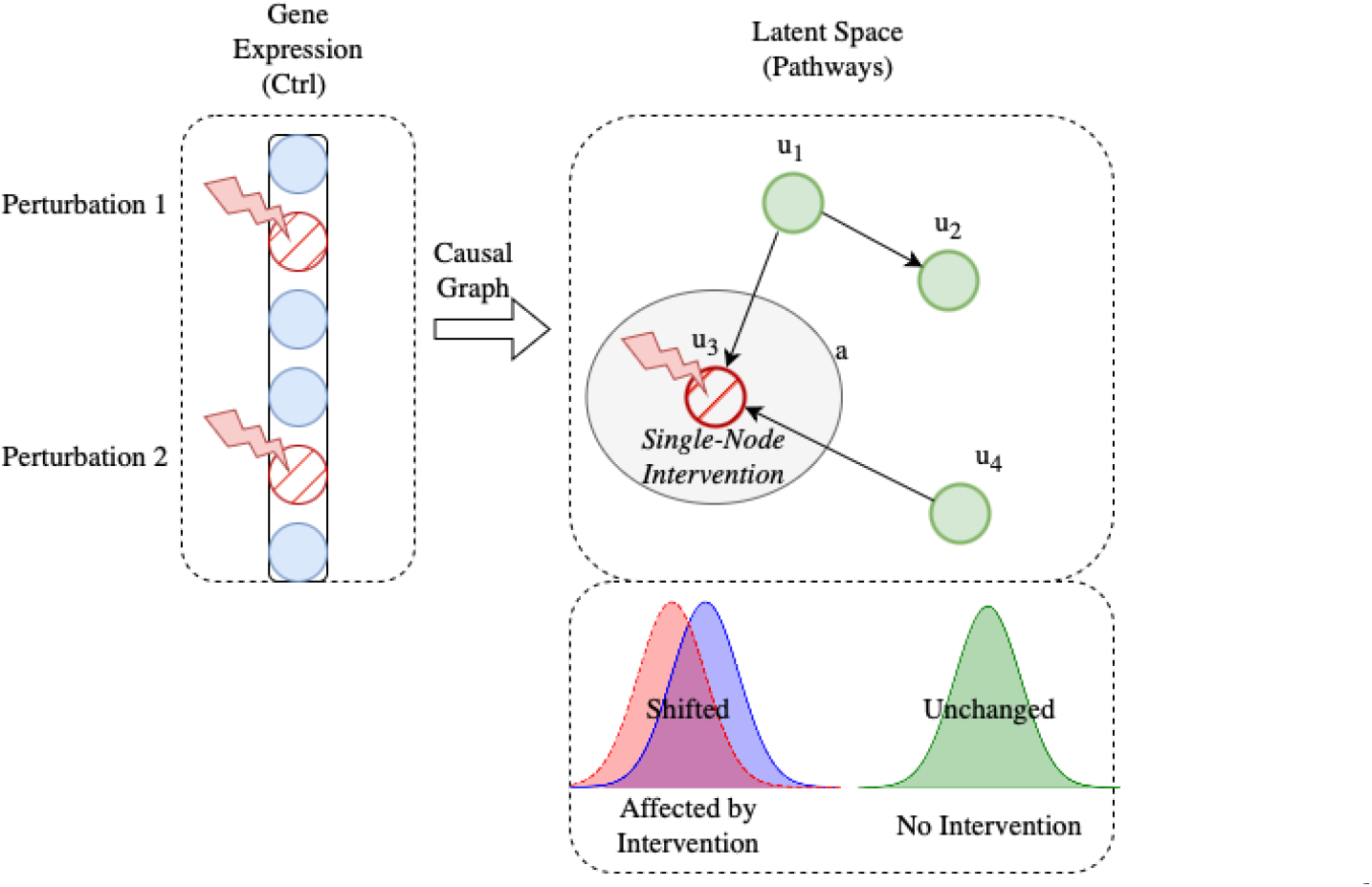
Illustration of an atomic (single-node) intervention in the latent space. An intervention in the observed gene expression space **x** causes a shift in a single latent pathway activity u_*i*_. This alters the conditional distribution *p*^*I*^ (u_*i*_ | Pa_*G*_ (u_*i*_)) while leaving unrelated latent variables unaffected, a key requirement for identifiability.

#### Requirement 1 (Acyclic Causal Structure)

We model the causal relationships among latent pathways using a directed acyclic graph (DAG), a standard structural assumption in causal discovery [19]. This assumption provides a well-defined causal ordering and excludes directed feedback cycles within the modeled structure.

#### Requirement 2 (Atomic, Single-Node Interventions)

Several causal representation learning methods establish identifiability guarantees under atomic interventions, in which each intervention directly modifies the causal mechanism of a single latent variable [9, 21]. In particular, Zhang et al. [9] consider single-target soft interventions and assume that every latent variable is targeted at least once. Under additional assumptions, this setting permits recovery of causal ancestry and the latent causal graph up to the relevant equivalence class.

#### Requirement 3 (Faithfulness)

Causal discovery commonly assumes faithfulness, meaning that conditional independencies in the observed distribution reflect graphical separation rather than coincidental cancellations of causal effects. Zhang et al. [9] introduce two additional conditions for identifying latent causal structures. First, *linear interventional faithfulness* requires that relevant intervention-induced distributional changes remain detectable under specified linear combinations of latent variables, enabling identification of ancestral relationships. Second, *adjacency faithfulness* excludes conditional independencies that could obscure direct causal connections under linear transformations, enabling recovery of the causal graph under the remaining identifiability assumptions.

Under these conditions, it is possible to provably recover the true latent causal graph up to a well-defined equivalence class. Following the work of Zhang et al. [9], their main theorem provides a formal guarantee for identifiability (Theorem 1):

**Theorem 1** (Full DAG Identifiability). *Assume an SCM with a latent DAG G and an invertible mixing function. If the set of interventions is atomic (single-node, targeting each latent variable at least once), and the assumptions of faithfulness hold, then the latent DAG G and the intervention assignments are identifiable up to permutation, scaling, and translation (the CD-equivalence class)*.

This reliance on an **atomic set of interventions**, where each latent variable is targeted at least once, is a cornerstone of modern CRL. It forms the basis for models applied to Perturb-seq data, as seen in recent methods [9, 10].

Importantly, we distinguish modeling requirements from theoretical identifiability assumptions and from empirical validation. The directed acyclic structure (Requirement 1) and the atomic, single-target intervention mapping (Requirement 2) are modeling requirements adopted in our architecture. Theoretical identifiability guarantees instead require intervention coverage, restrictions on the observation map, and the appropriate faithfulness conditions described above. Finally, whether the learned representations and perturbation effects are biologically meaningful is an *empirical* question. This distinction matters because Zhang et al. [9] prove identifiability under their specified generative assumptions; those guarantees do not automatically transfer to RAPTORGraph merely because it enforces a sparse intervention mapping and an acyclic graph. In particular, enforcing a single-node intervention in the learned representation does not prove that a biological perturbation directly changes only one latent causal mechanism: a genetic perturbation may have direct effects on multiple biological pathways, and assigning it to exactly one latent entry point is an architectural constraint rather than an experimentally verified property.

### The intervention spillover problem

While identifiability theory provides a strong theoretical foundation, a key practical challenge arises from the architectural choices in most CRL models. We term this the *intervention spillover problem*: the phenomenon where a targeted, single-variable intervention in the high-dimensional observation space (**x**) is transformed into a dense, multi-node signal in the lower-dimensional latent space (**u**).

This issue stems from a direct conflict with the atomic intervention requirement discussed above. An *Ideal Causal Encoder, f*^*∗*^, must satisfy this requirement by ensuring that a targeted intervention on a single gene isolates its distributional effect to the parameters of a corresponding single latent variable. Consequently, the expected change in the latent space would be perfectly sparse, manifesting as a clean, single-node perturbation rather than a dense signal.

#### Intervention spillover problem in current models

However, current interventional encoder networks *f*_*θ*_ are typically dense architectures like MLPs. These encoders possess a dense Jacobian matrix, meaning each component of the latent vector is a function of all input components. Consequently, a sparse change in the input **x** propagates through the network, and the model is thus forced to choose the ‘best’ single target from a dense signal that suggests multiple viable targets.

This architectural mismatch has several critical implications for the reliability of learned intervention targets. First, models that use a softmax function for target identification are forced into a “winner-take-all” decision on a dense signal, creating a risk of misattributing the primary causal effect. Furthermore, while the perturbed gene is known *a priori*, its corresponding target in the latent space is not, making the task of reliably discovering it from an entangled signal intractable. This reframes the primary modeling objective: the model should be designed not to explore for a latent target, but to accurately learn the magnitude of the intervention’s effect on the causal representation.

#### Spillover problem in genetic perturbation prediction

This phenomenon is particularly concerning in genetic perturbation experiments, as interventions are applied to individual genes in the high-dimensional expression space, yet their effects propagate through the cell’s regulatory network and manifest as coordinated changes across multiple (latent) biological processes. It is thus crucial to distinguish this intervention spillover, an artifact of dense encoders, from true biological correlation. High correlations between genes do not violate the single-node intervention requirement. Rather, dense encoders fail by transforming a sparse perturbation into a dense entry point in the latent space. By enforcing a sparse mapping for the intervention target, our proposed architecture satisfies identifiability while still modeling complex gene–gene correlations downstream via the dense components of the encoder and the learned causal graph.

### The dilemma of prior knowledge in causal discovery

Existing causal discovery models can be positioned on a spectrum from purely exploratory approaches like dVAE, which learns all components from data, to more confirmatory models like SENA, which leverages predefined biological pathways to enhance interpretability. While this confirmatory strategy aims to ground the model in known biology, it introduces a significant limitation stemming from poor knowledge transfer. Biological pathways are known to exhibit strong cell-type and cell-line specificity [22–25], meaning that mechanisms characterized in one cellular context often do not generalize to another.

## Results

To first establish the foundational accuracy of RAPTORGraph, we benchmarked its generative performance against four state-of-the-art models: SENA, scGPT, dVAE, and GEARS. We assessed two key aspects of prediction fidelity primarily using the single-cell perturbation dataset from Norman et al. [4]. We prioritized this dataset for the comprehensive comparison because the implementations of several baseline models have auxiliary files and preprocessing steps that are tightly coupled to it, making adaptation to other datasets non-trivial. However, to further demonstrate generalizability, we provide an additional evaluation on the Replogle2020 dataset, restricted to a comparison between RAPTORGraph and scGPT.

To validate our hyperparameter selection, we conducted a comprehensive ablation study on the *β*-VAE regularization parameter (*β*) and the DAGMA loss weights (*µ, λ*_1_). Our analysis reveals a critical trade-off between distribution matching and the preservation of biological heterogeneity. While higher regularization (*β ≥* 0.8) achieves competitive Maximum Mean Discrepancy (MMD) scores, it induces severe posterior collapse, with the variance of predicted control cell states dropping by an order of magnitude compared to the optimal setting (10^*−*4^ vs 10^*−*3^). Conversely, lower regularization (*β* = 0.1) preserves higher variance but results in unstable training and significantly higher MMD. We identified *β ≈* 0.4 as the optimal operating point, minimizing MMD while maintaining a latent KL divergence (*≈* 10^*−*3^) consistent with established baselines (dVAE and SENA), thereby ensuring the model captures meaningful cellular diversity rather than regressing to the population mean.

### Reconstruction capabilities

First, we measured the Mean Squared Error (MSE) between the ground truth and predicted profiles of control cells to evaluate how well each model reconstructs the basal, unperturbed cellular state. Second, we calculated the MMD between the distributions of ground truth and predicted profiles for double-gene perturbations to assess the model’s ability to capture the global transcriptional shift following a complex intervention.

The results in Table 1 highlight a crucial trade-off between reconstruction fidelity (MSE) and distributional accuracy (MMD). RAPTORGraph achieves a strong overall balance, demonstrating highly competitive performance across both reconstruction and generative capabilities.

**Table 1.** **Results using the Norman-CPA dataset**, averaged over 3 runs. Bold: best results; Italics: second best. Values are mean *±* standard deviation. *We excluded the GEARS model because it behaves in a fundamentally different way that is incompatible with our evaluation framework, making a fair and direct comparison impossible.

| Metric | SENA | scGPT | dVAE | GEARS | RAPTORGraph (ours) |
|---|---|---|---|---|---|
| MMD | 0.522 ± 0.003 | 0.176 ± 0.001 | **0.076 ± 0.000** | –* | *0.112 ± 0.001* |
| MSE | **0.043 ± 0.000** | 0.066 ± 0.000 | 0.062 ± 0.000 | –* | *0.045 ± 0.000* |

The evaluation on the independent Replogle2020 dataset, restricted to a direct comparison with scGPT, further demonstrates robust reconstruction fidelity and distributional accuracy on unseen biological data (Table 2).

**Table 2.** **Results using the Replogle2020 dataset**, averaged over 3 runs. Bold: best results. Values are mean *±* standard deviation.

| Metric | scGPT | RAPTORGraph (ours) |
|---|---|---|
| MMD | 0.138 ± 0.000 | **0.137 ± 0.000** |
| MSE | 0.069 ± 0.000 | **0.054 ± 0.000** |

### Prediction of non-additive genetic perturbations

This complex and biologically significant benchmark assesses a model’s ability to predict emergent effects that are not simple linear combinations of single-gene perturbations (Table 3). Specifically, we compared the models based on their ability to predict five distinct interaction types, using Precision@10 to quantify how effectively each model could identify true interactions within its top 10 predictions, a direct measure of its utility for experimental discovery.

**Table 3.** Comparison of non-additive genetic interaction prediction. Precision@10 scores for five models across distinct genetic interaction subtypes on the Norman et al. dataset. Higher is better. Values are mean *±* standard deviation over 3 runs.

| Metric | dVAE | SENA | GEARS | scGPT | RAPTORGraph (ours) |
|---|---|---|---|---|---|
| Synergy | 0.000 ± 0.000 | 0.233 ± 0.153 | *0.333* ± *0.115* | 0.100 ± 0.000 | **0.367** ± **0.058** |
| Suppression | 0.000 ± 0.000 | 0.167 ± 0.115 | *0.333* ± *0.115* | **0.400** ± **0.000** | 0.267 ± 0.153 |
| Neomorphism | *0.333* ± *0.153* | *0.333* ± *0.115* | 0.133 ± 0.058 | 0.100 ± 0.000 | **0.433** ± **0.058** |
| Redundancy | **0.933** ± **0.115** | 0.467 ± 0.058 | 0.567 ± 0.115 | 0.100 ± 0.000 | *0.800* ± *0.100* |
| Epistasis | *0.567* ± *0.153* | **0.800** ± **0.100** | 0.533 ± 0.115 | 0.400 ± 0.000 | 0.433 ± 0.115 |

We further investigated the contribution of OT to this performance. Our ablation studies reveal that OT preprocessing is a key driver of the model’s ability to predict complex non-additive interactions, particularly redundancy and epistasis, significantly outperforming a random pairing baseline. Importantly, this benefit comes with negligible computational overhead (*≈* 2.2% of total training time).

### Reverse perturbation analysis

While predicting genetic interactions demonstrates a model’s forward predictive power, a more profound test is its ability to reverse-engineer the genetic cause of an observed phenotype. To this end, we challenged a model to identify the specific genetic perturbation responsible for a given cellular gene expression profile. The analysis was performed on a controlled combinatorial space of 20 transcription factors from the Norman et al. [4] dataset. For each model, we generated a database of predicted expression profiles for all possible double-gene perturbations within this subset. We then used the ground truth mean expression profiles from experimentally observed double perturbations as queries, ranking the database entries by similarity to identify the most likely causal perturbation.

We first evaluated the models on the highly stringent task of identifying the exact causal gene combination, quantified by the Correct (Exact Match) Hit Rate @ K (S1 Fig). As shown there, all models achieve modest hit rates, underscoring the profound difficulty of this reverse-engineering challenge. However, a more practical measure of a model’s utility is its ability to identify biologically influential genes. We assessed this using the Relevant (Gene Overlap) Hit Rate @ K, which credits a model for retrieving perturbations containing at least one of the correct genes (Figure 2 and Table 4). RAPTORGraph demonstrates competitive performance, consistently ranking among the top models. This highlights that while pinpointing exact higher-order interactions is an ongoing challenge, RAPTORGraph is highly effective at identifying the key genetic players driving a cellular response, making it a valuable tool for hypothesis generation and experimental design.

**Table 4.** **Hit Rate @ K (relevant)** for the reverse perturbation task. The metric measures the fraction of queries where at least one correct gene was identified in the top K predictions. Higher values are better. Values are mean *±* standard deviation over 3 runs.

| *k* | dVAE | SENA | GEARS | scGPT | RAPTORGraph (ours) |
|---|---|---|---|---|---|
| 1 | 0.333 ± 0.115 | 0.244 ± 0.019 | **0.344** ± **0.051** | 0.233 ± 0.000 | 0.294 ± 0.059 |
| 3 | 0.556 ± 0.051 | 0.444 ± 0.069 | 0.578 ± 0.051 | 0.533 ± 0.000 | **0.608** ± **0.122** |
| 5 | 0.733 ± 0.088 | 0.600 ± 0.058 | 0.722 ± 0.077 | 0.700 ± 0.000 | **0.784** ± **0.090** |

**Fig 2.**
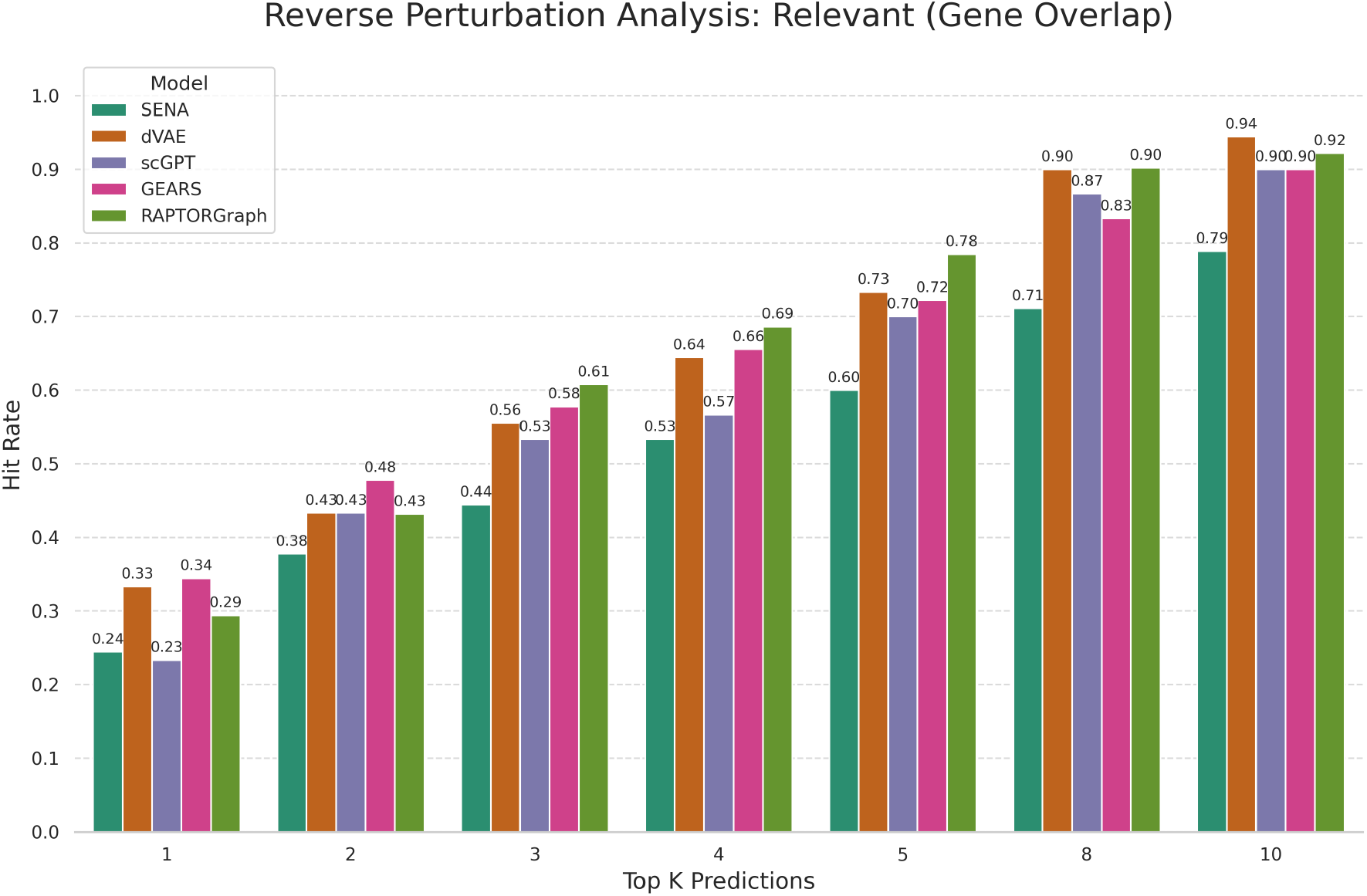
**Relevant (Gene Overlap) Hit Rate @ K** for the reverse perturbation task. The metric measures the fraction of queries where at least one correct gene was identified in the top K predictions. Higher is better. Values are mean *±* standard deviation over 3 runs.

**RAPTORGraph learns biologically meaningful causal latent factors**. To confirm that RAPTORGraph learns biologically meaningful structures rather than arbitrary representations, we examined the functional identity of the latent factors using gene set enrichment analysis (GSEA). For each learned latent factor, we identified the single-gene interventions that induce the greatest change in that factor and tested the resulting gene signatures against the “Hallmark” gene sets from the Human Molecular Signatures Database (MSigDB) [26]. It is important to distinguish these GSEA-derived pathways, which are well-described biological processes from curated databases, from the learned causal meta-pathways, which are latent factors associated with single interventions; our goal is to assess whether the learned meta-pathways encompass relevant and related biological processes. Figure 3 visualizes the Normalized Enrichment Scores (NES) for the top enriched pathways across the learned factors.

**Fig 3.**
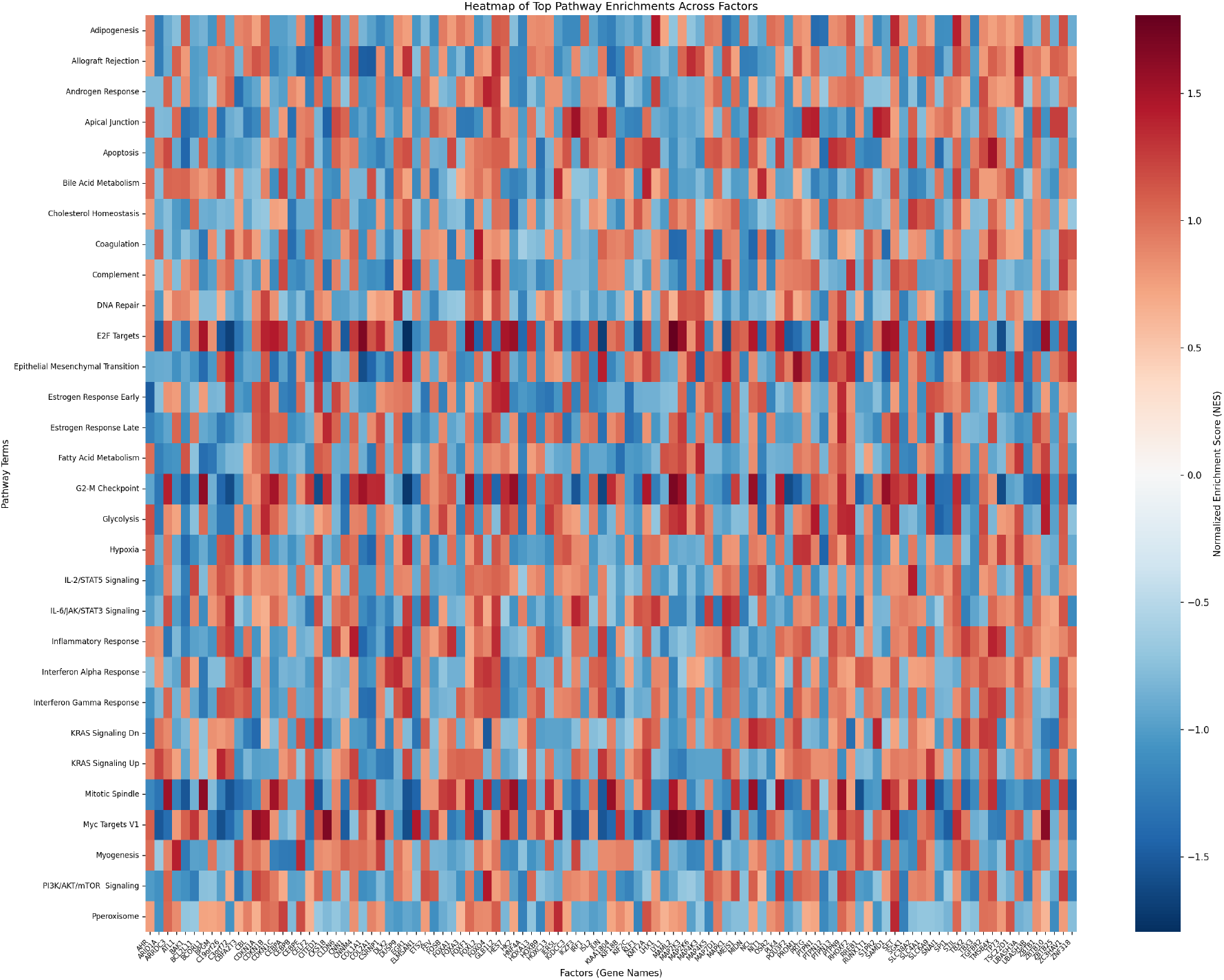
Normalized Enrichment Scores (NES) of the top enriched Hallmark pathways across the learned latent factors of RAPTORGraph.

The GSEA analysis reveals that RAPTORGraph captures distinct and biologically accurate programs, verified against the known functions of their corresponding gene perturbations:

1. **Differentiation-induced arrest:** The latent factor linked to *EGR1* displays massive negative enrichment for “E2F Targets” and “G2-M Checkpoint”. EGR1 is a known tumor suppressor in K562 leukemia cells that drives differentiation along the megakaryocytic lineage, a process that necessitates a complete exit from the cell cycle [27]; RAPTORGraph correctly identifies this potent anti-proliferative program.
2. **Stress-induced growth arrest:** The factor linked to the AP-1 subunit *JUN* shows strong negative enrichment for “E2F Targets” and “G2-M Checkpoint”. While often associated with inflammation, c-Jun is a central stress-response mediator; in K562 cells its activation drives a termination of cell division required for stress adaptation or differentiation [28, 29].
3. **p53-like tumor suppression:** The *TP73* -associated factor is negatively enriched for “G2-M Checkpoint”. *TP73* is a functional homolog of *TP53* that triggers downstream checkpoint signaling to arrest the cell cycle and prevent genomic instability [30].

These findings confirm that the nodes in our learned DAG represent specific biological mechanisms and that the edges learned by the DAGMA module can be interpreted as causal regulatory links between verified biological processes.

## Discussion

RAPTORGraph was designed to resolve two failure modes that limit existing perturbation-prediction models: intervention spillover, where dense encoders create entangled latent signals that confound causal discovery, and mean collapse, where the absence of one-to-one cell pairings erodes cellular diversity. Our results indicate that a preconditioned GraphPathway encoder can enforce the sparse, single-node interventions that causal identifiability theory requires, while a DAGMA module subsequently learns the legitimate biological causal relationships among the resulting sparse pathway activities. Because the model does not rely on predefined pathway annotations, it avoids the confirmatory bias of methods such as SENA, which must explain the data through preexisting pathways and cannot readily accommodate novel signals outside those annotations.

A recurring theme in our findings is the trade-off between reconstruction fidelity and distributional accuracy. RAPTORGraph achieves a competitive balance, and the optimal-transport preprocessing step proves important for predicting complex, non-additive interactions, particularly redundancy and epistasis, at negligible computational cost. The reverse-perturbation analysis further shows that the model can identify the key genetic drivers of an observed phenotype, supporting its use for hypothesis generation.

Several limitations remain. Our comprehensive baseline comparison was performed primarily on the Norman-CPA dataset because several baselines are tightly coupled to its preprocessing; generalizability evidence on the Replogle2020 dataset is currently limited to a comparison with scGPT. The exact-match hit rates in the reverse-perturbation task remain modest for all models, indicating that pinpointing precise higher-order combinations is an open challenge. Future work will focus on deploying RAPTORGraph for the hypothesis-driven discovery of novel drug targets and exploring its utility in uncovering cell-type-specific pathway mechanisms.

## Conclusion

Developing interpretable *in silico* models is crucial for therapeutic discovery, yet current approaches face two key challenges: intervention spillover, where dense encoders create entangled latent signals that confound causal discovery, and mean collapse, where the absence of one-to-one cell pairings erodes cellular diversity. We introduce RAPTORGraph, a framework that directly resolves these issues. To counteract intervention spillover, it combines a preconditioned GraphPathway layer, enforcing the sparse mappings required for clean single-node interventions, with a DAGMA layer for robust DAG discovery. This architectural design eliminates the artifactual entanglement of input genes with latent pathways, while the subsequent DAG module explicitly learns the legitimate biological causal relationships and correlations between the resulting sparse pathway activities. To prevent mean collapse, RAPTORGraph globally pairs control and perturbed populations through OT. RAPTORGraph not only achieves a strong balance between reconstruction fidelity and distributional accuracy relative to state-of-the-art methods, but also shows the ability to predict complex, non-additive genetic interactions, and can reverse-engineer genetic drivers to generate testable hypotheses. RAPTORGraph bridges predictive power and mechanistic insight, enabling not only prediction of a perturbation’s effect but also understanding of why. Future work will focus on deploying RAPTORGraph for the hypothesis-driven discovery of novel drug targets and exploring its utility in uncovering cell-type-specific pathway mechanisms.

## Materials and methods

### GraphPathway encoder for deconfounding through preconditioning

To address the Intervention Spillover Problem and enforce the theoretical requirement for single-causal-factor interventions, we introduce a novel encoder architecture, the GraphPathway Encoder, which features a deconfounding approach through preconditioning. This approach requires a specific ordering of the input data. Let *k* be the number of known perturbed genes in the experiment. Without loss of generality, we assume the input gene vector **x** *∈* R^*n*^ is sorted such that the first *k* genes are those perturbed, followed by the remaining *n − k* unperturbed genes.

**Pre-conditioning module (Figure 4-A)**. Given this sorted input, we define our model with *d* learned causal latents (meta-pathways), where we require that the number of learned meta-pathways is at least the number of perturbed genes (*d ≥ k*). We then structure the encoder’s primary weight matrix ***W*** as a block matrix:

**Fig 4.**
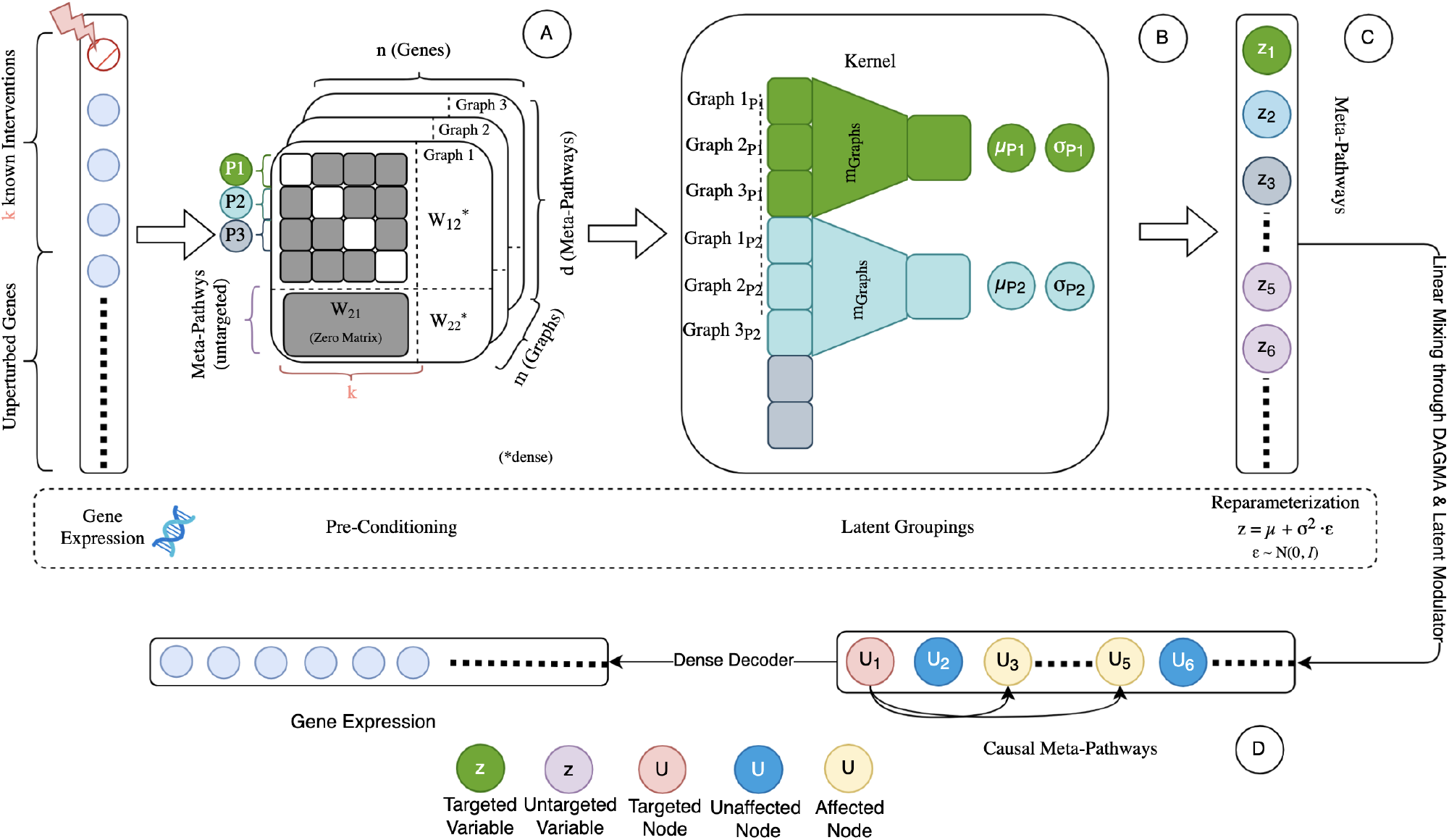
The GraphPathway Encoder architecture of RAPTORGraph. **A**: Gene expression of control cells (**z**_obs_) is fed as input to the preconditioned linear layer, which enforces a sparse, one-to-one mapping for known perturbed genes to explicitly prevent intervention spillover. **B**: Complex dependencies are captured by learning multiple parallel graph representations, which are then aggregated to model non-linear interactions. **C**: A standard VAE reparameterization head generates the stochastic latent variables (**z**). **D**: A Latent Modulator takes the perturbation label as input and applies atomic perturbations (∆) to the independent latents (**z**), shifting the distribution from the observed latent state (**z**_obs_) to the interventional latent state (**z**_int_). Both **z**_obs_ and **z**_int_ are then transformed by the DAGMA layer through the learned DAG via linear mixing to yield the final causal states (**u**), which are reconstructed into the predicted control 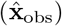 and perturbed 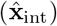 gene expression profiles by a dense decoder.

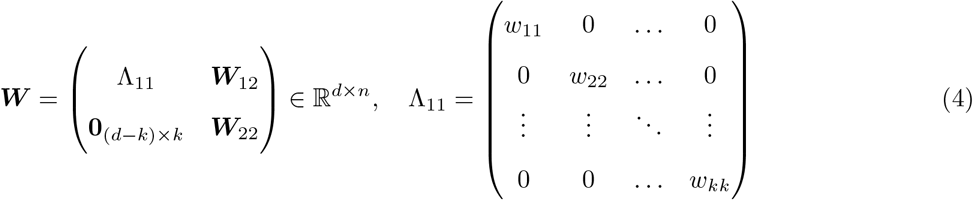

where the top-left submatrix Λ_11_ *∈* R^*k×k*^ is a learnable, strictly diagonal matrix.

This diagonal structure enforces a direct, one-to-one mapping, ensuring each of the *k* perturbed genes directly influences only one of the first *k* learned meta-pathways. The bottom-left zero block, **0**_(*d−k*)*×k*_, is critical for preventing intervention spillover by explicitly blocking the influence of perturbed genes on the remaining learned meta-pathways. The other submatrices, ***W***_12_ and ***W***_22_, are learnable and dense, allowing the model flexibility to discover complex relationships between unperturbed genes and all learned meta-pathways. Crucially, this preconditioned structure allows the intervention mask ***m*** to be predefined and fixed rather than learned, directly satisfying the theoretical requirements for identifiability.

#### Latent grouping via an interaction block (Figure 4-B)

To model the complex non-linear dynamics of biological systems, our architecture extends this preconditioning approach by learning multiple distinct subgraph representations for each (exogenous) latent factor. An intermediate *Interaction Block* then processes these subgraph activations, allowing the model to capture higher-order dependencies by learning unique interaction patterns among the subgraphs within each learned latent factor.

#### Exogeneous latent factor inference via VAE (Figure 4-C)

Next, an aggregation layer combines these processed features to compute the parameters (*µ*_**z**_ and *σ*_**z**_) for the (exogenous) latent distributions. Thus, the final stochastic representations of the latent factors are then obtained using the reparameterization trick **z** = *µ*_**z**_ + *σ*_**z**_ *⊙ ϵ*; *ϵ ∼ N* (**0, *I***). Subsequently, a latent modulator predicts a conditional shift for the targeted latent variables, strictly modifying only the specific latent variable associated with the perturbation while preserving the independence of the others.

#### Causal latent factor (causal meta-pathway) inference via DAGMA (Figure 4-D)

To capture the downstream biological consequences, a DAGMA layer propagates these atomic interventions through a learned directed acyclic graph, transforming the independent noise terms into causally dependent pathway activities. Finally, a non-linear dense decoder maps these propagated causal states back to the high-dimensional gene expression space to predict the final post-perturbation phenotype.

### Deep causal graph learning with DAGMA

The GraphPathway Encoder with preconditioning fundamentally alters the causal discovery task. By fixing the mapping from known interventions to specific latent variables, it eliminates the need for permutation-based discovery methods that assume a dense, entangled encoder. However, this fixed ordering makes simple acyclicity constraints, such as enforcing an upper-triangular adjacency matrix, unsuitable.

These methods impose a strict and arbitrary causal hierarchy, as the influence of one latent variable on another is entirely restricted by their ordering (e.g., latent *i* can only influence latent *j* if *i < j*), which would conflict with the true biological relationships we aim to discover. For this, we adapt the DAGMA framework [31] for use in a deep learning context.

**DAGMA** reformulates the combinatorial search for a DAG into a continuous optimization problem, following the continuous acyclicity-constrained paradigm of NOTEARS [32]. This is achieved by minimizing a score function that measures data fit, subject to a differentiable constraint that is satisfied only if the graph is acyclic. The DAGMA acyclicity constraint, *h*_*s*_(***A***), is based on a log-determinant function:

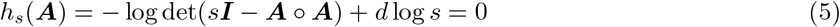

where ***A*** *∈* R^*d×d*^ is the weighted adjacency matrix of the latent graph, *s >* 0 is a scaling scalar, and the condition holds if and only if ***A*** represents a DAG. The optimization then solves a sequence of unconstrained problems that jointly minimize a data-fit score and the acyclicity penalty.

#### DAGMA implementation

We implement a modified DAGMA approach designed to learn the invariant, baseline causal structure of the biological system. To ensure stability and manage the trade-off between reconstruction fidelity and acyclicity, we introduce two critical regularizations. First, we employ a path-following schedule where the score weight *µ* is gradually decreased (see Eq. (7)), allowing the model to prioritize feature learning early in training before strictly enforcing the graph constraint. Second, we isolate gradient flow by computing *L*_DAGMA_ on detached latent representations. This ensures the acyclicity constraint optimizes the graph structure without distorting the encoder’s feature extraction.

### Training objective

The model is trained end-to-end by minimizing a composite objective function, *L*_total_, that integrates four distinct components, each tailored to a specific aspect of the learning problem. The overall objective is a weighted sum of these individual losses:

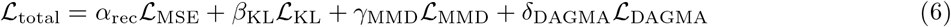

where *α*_rec_, *β*_KL_, *γ*_MMD_, and *δ*_DAGMA_ are scalar hyperparameters that balance the contribution of each term.

#### Reconstruction Loss (*L*_MSE_)

For observational data, the model is trained to faithfully reconstruct the input gene expression profiles. This is measured by the MSE between the decoder’s output and the original unperturbed input, ensuring the model learns a high-fidelity representation of the basal cellular state.

#### KL Divergence (*L*_KL_)

As a standard component of a VAE, a Kullback-Leibler divergence term is included to regularize the latent space. It encourages the learned posterior distribution over the latent variables, *q*(**z**|**x**), to be close to a prior distribution, *p*(**z**), which is typically a standard normal distribution. This promotes a smooth and well-structured latent space.

#### Interventional Prediction Loss (*L*_MMD_)

For interventional data, the model’s ability to predict out-of-distribution outcomes is evaluated using the MMD [33]. This loss measures the distributional distance between the model-predicted cell states following a perturbation and the ground-truth distribution of experimentally observed cells for that same perturbation.

#### Causal Graph Loss (*L*_DAGMA_)

The DAGMA loss is computed on the latent representations of observational data to learn the invariant causal graph structure between the learned meta-pathways. This objective integrates three key components: a data-fit score, an L1 penalty to encourage sparsity, and the differentiable acyclicity constraint. The loss is defined as:

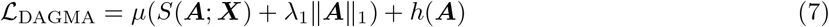

where *S*(***A***; ***X***) is a score, ∥***A***∥_1_ is the L1 norm of the adjacency matrix, and *h*(***A***) is the acyclicity penalty. The hyperparameters *µ* and *λ*_1_ control the weighting of the score and sparsity terms, respectively.

### Optimal transport for deterministic interventional learning

A fundamental challenge in learning from single-cell perturbation data is the absence of a direct one-to-one mapping between control and perturbed cell populations, as it is impossible to observe the same cell before and after the intervention. This makes learning the treatment effect a fundamental counterfactual problem, where the central challenge is to identify a general transformation that maps the control distribution to the perturbed distribution in the absence of paired examples. To address this counterfactual gap, many models create training pairs by randomly matching control and interventional samples. While computationally efficient, this random pairing is a primary cause of what we term *mean collapse*: a process where the model learns the average response, collapsing the distribution towards its mean and eroding cellular diversity (Fig. 5).

**Fig 5.**
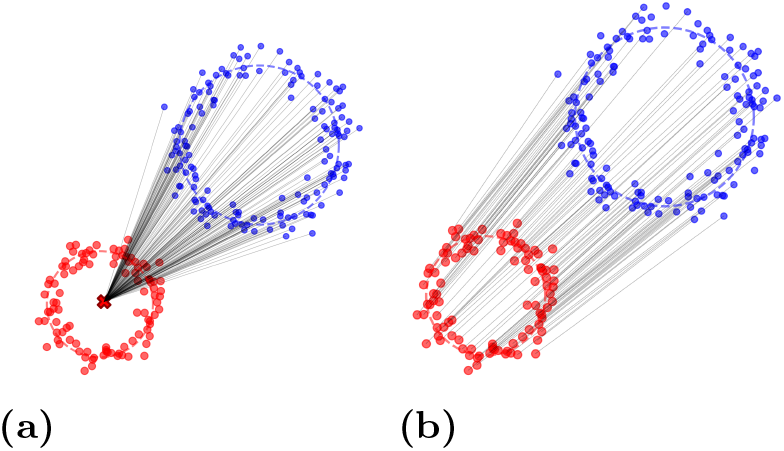
Illustrative example of mean collapse problem. (a) Random pairing collapses the source distribution towards the target mean. (b) The OT preserves the underlying population structure.

To address these issues, we employ a Wasserstein-based OT [34, 35] framework to globally pair control cells with perturbed cells under unique conditions (Fig. 5b). Although several recent methods address unpaired single-cell perturbation data [17, 36], we adopt OT as an offline pairing step for simplicity while avoiding mean collapse.

### Evaluation setup and metrics

To ensure a consistent and fair benchmark across all models, we employ a suite of metrics targeting different aspects of model capability. We use the MSE to evaluate reconstruction of basal (control) cell states, the MMD to evaluate the accuracy of out-of-distribution predictions for perturbed states, Precision@10 to evaluate non-additive genetic interactions, and Hit Rate @ K to evaluate reverse-engineering of genetic causes. The calculation of all metrics was standardized across the benchmarked tools.

#### Training details

RAPTORGraph is implemented in PyTorch Lightning and trained with the AdamW optimizer at a learning rate of 10^*−*4^ and weight decay of 1.3 *×* 10^*−*4^. We use a batch size of 256 and train for 100 epochs on a single GPU. The GraphPathway encoder uses a hidden dimension of 904, eight pathway graphs, and three interaction blocks with SiLU activations and residual connections; the decoder is a two-layer MLP that maps the latent space back to the gene-expression space. The number of latent factors is set to 1.04*×* the number of perturbed genes. The composite objective (Eq. (6)) combines the reconstruction term (weight *α*_rec_ = 1) with the MMD term weighted by *γ*_MMD_ = 0.197, the KL term weighted by *β*_KL_ = 0.447, and the graph term weighted by *δ*_DAGMA_ = 1; the DAGMA loss itself uses score weight *µ* = 0.27 and sparsity weight *λ*_1_ = 0.052. All reported results are averaged over three random seeds (41, 42, 43).

#### Mean Squared Error (MSE)

The MSE measures each model’s ability to faithfully reconstruct the gene expression profile of control (unperturbed) cells; a lower value indicates better performance. To avoid trivial memorization, the set of control cells is randomly split into two non-overlapping halves: an input set and a ground-truth set. The model reconstructs cells from the input set, and the MSE is computed between the reconstructions and the corresponding ground-truth cells, averaged over batches. For a batch of *i* cells with *n* genes, the MSE is

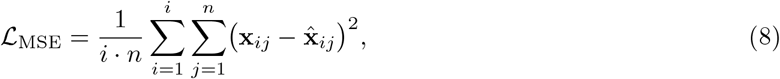

where **x**_*ij*_ is the expression level of gene *j* in true control cell *i*, and 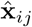 is the corresponding reconstructed value.

#### Maximum Mean Discrepancy (MMD)

The MMD [33] measures the distance between two probability distributions based on the distance between their mean embeddings in a reproducing kernel Hilbert space. It quantifies the dissimilarity between the distribution of model-predicted gene expression profiles and the distribution of true, experimentally observed profiles for a given perturbation. We use a Gaussian (radial basis function) kernel

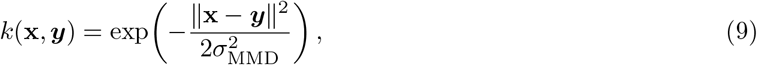

where *σ*_MMD_ is the bandwidth hyperparameter. The squared MMD between the true distribution *p*_data_ and the model’s distribution *p*_model_ is, in population form,

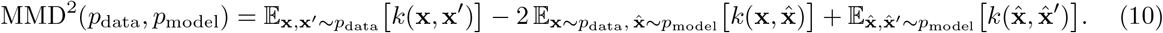

In practice we use a multi-kernel (multi-window) estimate, i.e. a weighted combination of Gaussian kernels at several bandwidths, and the unbiased empirical estimator on batches; we refer to this estimator as MMD throughout. The MMD is calculated per-condition and then averaged to provide an overall measure of distribution similarity.

#### Precision@10 for non-additive genetic interactions

To evaluate the models’ ability to predict non-additive effects, we use the Precision@10 metric inspired by the GEARS study [5], with a naive additive baseline. For each double-gene perturbation, we compute an effect vector ∆ as the mean perturbed profile minus the mean control profile, for both ground-truth data and model predictions. The naive additive baseline assumes the double-perturbation effect is the sum of the individual single-perturbation effects, ∆(*A* + *B*)_naive_ = ∆_*A*_ + ∆_*B*_. Comparing the true or predicted combined effect ∆(*A* + *B*) against this baseline yields five interaction scores:

- *Synergy* and *Suppression*: Synergy Ratio = 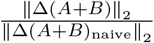(magnitude ratio);
- *Neomorphism*: Neomorphism Score = 1 *−* cos (∆(*A* + *B*), ∆(*A* + *B*)_naive_) (change in direction);
- *Redundancy* : Redundancy Score = min cos (∆(*A* + *B*), ∆_*A*_), cos (∆(*A* + *B*), ∆_*B*_) (functional overlap);
- *Epistasis*: Epistasis Score = cos |(∆(*A* + *B*), ∆_*A*_) *−* cos (∆(*A* + *B*), ∆_*B*_)| (dominance).

For each interaction type, perturbations are ranked by their predicted score and the top 10 are selected. These are compared against a ground-truth set of interactors, defined as those whose true score falls in the top 25% (or, for suppression, the bottom 25%) of that category. Precision@10 is the fraction of the top-10 predictions that are true interactors; higher is better.

#### Hit Rate @ K for reverse perturbation

To assess each model’s ability to reverse-engineer the genetic cause of an observed phenotype, we query the model with a ground-truth mean expression profile 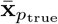of a double perturbation and rank the database of predicted profiles by Euclidean distance to the query. We report two variants of Hit Rate @ K. *Correct (Exact Match) Hit Rate @ K* is the fraction of queries for which the exact true perturbation is within the top K predictions. *Relevant (Gene Overlap) Hit Rate @ K* is the fraction of queries for which at least one of the true perturbation’s genes is present in the top K predictions, i.e. genes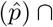 genes(*p*_*q*_) ≠ *∅* for some prediction 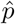 in the top K.

We used the Perturb-seq single-cell RNA-seq dataset curated by Norman et al. [4] as processed in the CPA study [16] and refer to it as the Norman-CPA dataset. It profiles 105 gene perturbations in K562 cells, including both single and double perturbations. In total, 284 conditions across *∼*108,000 cells were measured, of which 131 represent unique gene–gene combinations and the remainder single perturbations or controls, respectively. We utilize the entire set of single-perturbations during training and evaluate the subsequent methods on the double-perturbations.

Additionally, we used the CRISPRi single-cell RNA-seq dataset curated by Replogle et al. [13] as processed in the PerturBase study [37] and refer to it as the Replogle2020 dataset. This dataset profiles 82 gene perturbations in K562 cells, including both single (44) and double (37) perturbations and control. The 82 conditions were measured across *∼*22,740 cells.

## Acknowledgments

The authors acknowledge the financial support by the Ministry of Science and Culture of Lower Saxony through the *zukunft*.*niedersachsen* program of the Volkswagen Foundation via the CAIMed (Lower Saxony Center for Artificial Intelligence and Causal Methods in Medicine) project under project number ZN4257, and by the German Federal Ministry of Education and Research (BMBF) in the framework of the P4D (Personalisierte, prädiktive, präzise und präventive Medizin zur Verbesserung der Früherkennung, Diagnostik, Therapie und Prävention depressiver Erkrankungen) project under project number 01EK2204F. MH: Ramon y Cajal fellowship (RYC2021-033127-I) and DL2CURE project (PID2023-151980OB-I00) funded by MCIN/AEI/10.13039/501100011033 and “NextGenerationEU”/PRTR; Palatchi Foundation; AECC Proyectos generales 2025 (PRYGN259200HERN). Views expressed herein are solely those of the author(s) and do not necessarily reflect the views of the German Federal Government, the State of Lower Saxony, nor the granting authorities.

## Data Availability

The source code of RAPTORGraph is available at https://github.com/sXperfect/RAPTORGraph. The Norman-CPA Perturb-seq dataset is available from the original Norman et al. study as processed in the CPA study [4, 16]; the Replogle2020 CRISPRi dataset is available from Replogle et al. as processed in the PerturBase study [13, 37].

## Supporting information

- **S1 Fig. Correct (Exact Match) Hit Rate @ K** for the reverse perturbation task. The metric measures the fraction of queries where the exact causal gene combination was identified in the top K predictions. Higher is better. Values are mean *±* standard deviation over 3 runs.
- **S1 Table**. Full hyperparameter sweep and ablation results for the *β*-VAE regularization parameter and DAGMA loss weights.

